# High concentration and yield production of mannose from açaí (*Euterpe oleracea*) seeds via diluted-acid and mannanase-catalyzed hydrolysis

**DOI:** 10.1101/513168

**Authors:** Alvaro Ferreira Monteiro, Ingrid Santos Miguez, João Pedro R. Barros Silva, Ayla Santana Silva

**Affiliations:** Biocatalysis Laboratory, National Institute of Technology, Ministry of Science, Technology, Innovation and Communication, 20081-312, RJ, Brazil; Federal University of Rio de Janeiro, Department of Biochemistry, 21941-909, RJ, Brazil

**Keywords:** Açaí, seed, mannan, diluted-acid hydrolysis, mannanases, mannose

## Abstract

The açaí berry’s seed corresponds to 85–95% of the fruit’s weight and represents ~1.1 million tons of residue yearly accumulated in the Amazon region. This study confirmed that mannan is the major component of mature seeds, corresponding to 80% of the seed’s total carbohydrates and about 50% of its dry weight. To convert this high mannan content into mannose, a sequential process of diluted acid and enzymatic hydrolysis was evaluated. Diluted-H_2_SO_4_ hydrolysis (3%-acid, 60-min, 121°C) resulted in a 30% mannan hydrolysis yield and 41.7 g/L of mannose. Because ~70% mannan remained in the seed, a mannanase-catalyzed hydrolysis was sequentially performed with 2–20% seed concentration, reaching 146.3 g/L of mannose and a 96.8% yield with 20% solids. As far as we know, this is the highest reported concentration of mannose produced from a residue. Thus, this work provides fundamental data for achieving high concentrations and yields of mannose from açaí seeds.

**Highlights:**

- Mannan was confirmed as the major component (~50%) of açaí seeds.
- Diluted-H_2_SO_4_ hydrolysis had a limited effect on mannan conversion into mannose.
- Enzymatic hydrolysis was sequentially performed with a high seed concentration.
- Mannan was efficiently hydrolyzed by mannanases, producing a 96.8% yield.
- Mannose production of 146.3 g/L was obtained with mannanase-catalyzed hydrolysis.

**Graphical abstract:** 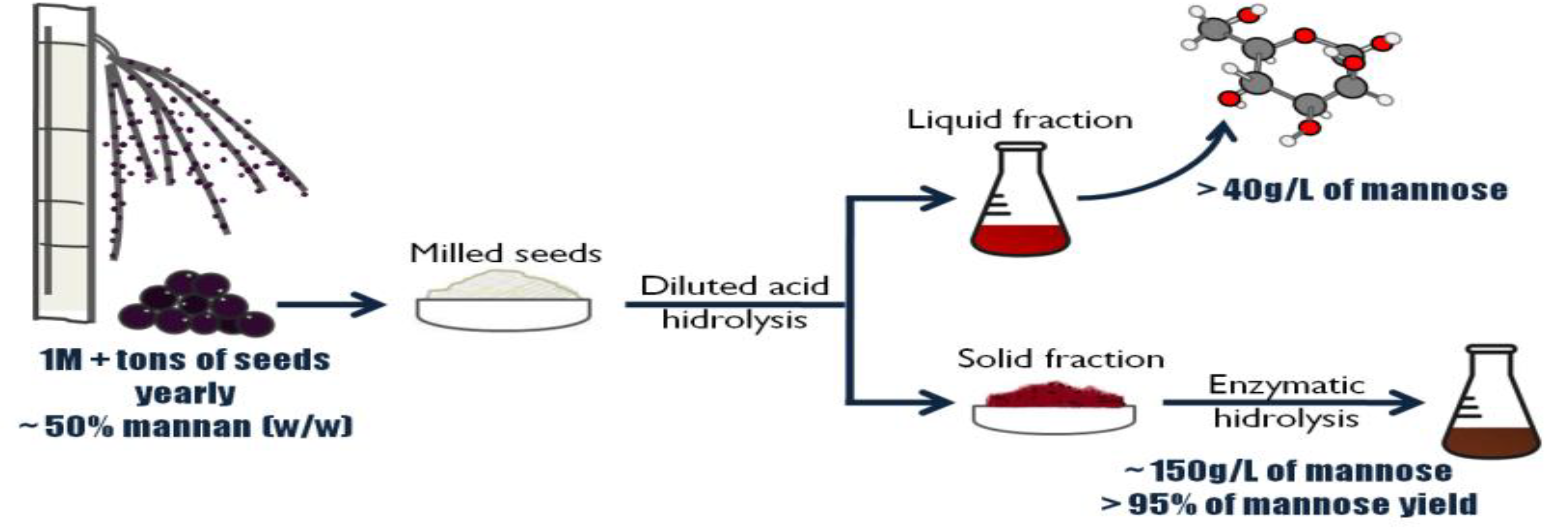

## 1. Introduction

The *Euterpe oleracea* palm plant—otherwise known as the açaí palm—is a widely distributed plant in northern South America, with a large percentage of its population found in the delta region of the Amazonian river (Yamaguchi et al., 2015). In the past 15 years, the commercialization of its fruit—the açaí berries—has experienced an economic boom, with a sizable increase seen in national and international markets, such as the United States, Japan, and Europe (Fioravanti, 2013; Nogueira et al., 2013). The main açaí producer in the world is the Brazilian state of Pará, representing more than 95% of Brazil’s production. This state produced more than 1,274,000 tons of açaí berries in 2017 (http://sidra.ibge.gov.br and http://www.sedap.pa.gov.br), and governmental incentives were recently created (Pró-Açaí Program) to increase the production by 360,000 tons by 2024.

The commercialized product—the açaí pulp—represents only 5–15% of the fruit’s weight, whereas the açaí seed accounts for the other 85–95% (Pessoa et al., 2010; Pompeu et al., 2009). The rapid increase of açaí commercialization has generated an enormous amount of açaí seeds as a residue of the extraction process, which is estimated at more than 1,000,000 tons deposited yearly in the Amazon region, with the prospect of more in the coming years. Today, only a small amount of the seeds is utilized for animal feed, plantations, or home gardens and crafts, and very few appropriate disposal methods currently exist, resulting in an acute environmental and urban problem (Fioravanti, 2013). It is of great environmental and economic interest to avoid waste production and simultaneously find new applications for açaí seeds, thus adding value to the productive chain and promoting local and social development.

To extract the highest possible value from this residue and determine the appropriate applications, it is of the utmost importance to know its composition, but few systematic studies have been carried out with açaí seeds. Previous works have reported that açaí seeds contain high amounts of carbohydrates (~70%), with cellulose being the main polysaccharide (Altman, 1956; Oliveira et al., 2015, 2013; Rodríguez-Zúñiga et al., 2008; Wycoff et al., 2015). In contrast, Rambo et al. have shown that 53.8% of the seed is composed of mannan, a polymer of mannose (Rambo et al., 2015). Therefore, the confirmation of the actual composition of the açaí seed and studying the processing methods to release its sugars is extremely relevant because the possibly unprecedented mannose content renders açaí seeds as a valuable and unexplored material.

One alternative to explore the açaí seed’s potential is to develop mild methods that efficiently release sugars from the material. Sugars derived from hemicellulose can be directly released from plant cell walls by applying different methods, such as acid, thermal, enzymatic hydrolysis, or microbial fermentation (Hu et al., 2016). Commonly, diluted inorganic acids, including sulfuric and hydrochloride acid, have been employed to hydrolyze the xylan-containing hemicelluloses of some lignocellulosic biomasses with more than 90% efficiency, resulting in a liquid fraction rich in free xylose and a solid residue rich in cellulose and lignin. Nevertheless, some studies have indicated that mannan-containing hemicelluloses, such as softwoods, may be less prone to the action of diluted sulfuric acid in a single step hydrolysis (Lim and Lee, 2013). On the other hand, mannan-degrading enzymes could also be applied for the release of free mannose from mannan-rich residues (Kusakabe et al., 1987; Srivastava and Kapoor, 2017; van Zyl et al., 2010), which could also be combined in a sequential step after a mild diluted acid hydrolysis. The enzymatic process could permit to work with mild conditions that are less damaging to the environment, besides improving the yield using simple protocols (Jiang et al., 2018). Up to now, there are no reports of studies aiming to release mannose from açaí seeds, which is a sugar with a high potential to be a functional ingredient and that exhibits biological functions of great interest in the cosmetic, pharmaceutical, and food industries (Ariandi et al., 2015; Otieno and Ahring, 2012; Rungrassamee et al., 2014).

For example, mannose can be easily reduced to mannitol—a specialty chemical with a wide variety of uses in the pharmaceutical industry—in a process with a 90% yield (Mishra and Hwang, 2013). However, because of the lack of abundant and low-cost sources of mannose, the conventional industrial processes for mannitol production are based on the chemical hydrogenation of fructose or inverted sucrose, which produce low yields of about 25% and 50%, respectively (Makkee et al., 1985).

Considering that açaí seeds can be a potential rich source of mannan, its high abundance in Brazil, and the limited current knowledge of mannan depolymerization, the aim of the current study was to confirm the carbohydrate composition of açaí seeds and to evaluate acidic-and enzymatic-catalyzed strategies to maximize mannose production.

## 2. Material and Methods

### 2.1. Source of materials

Açaí seed samples were kindly donated by the company Açaí Amazonas. Mannanase BGM “Amano” 10 was kindly provided by Amano Enzyme Inc. (Japan). All other chemicals were purchased from commercial sources and used without any further purification.

### 2.2. Characterization of açaí seeds

Two lots of açaí seeds were characterized in the current study. Samples from lot 1 were received as shown in Figure 1a and were noted as “whole seeds” (Fig. 1a), while lot 2 contained samples already milled (Fig. 1d). For characterization, the whole seeds samples from lot 1 were processed with a knife mill as received. Alternatively, for lot 1 samples, the external fiber layers (Fig. 1c) were manually separated from the inner core stones (Fig. 1b), and the two fractions were milled separately for further chemical characterization. In parallel, the masses of 35 whole seed samples were measured using an analytical balance (0.001 precision). Subsequently, the external fiber layers were manually removed from the core stones and then both were weighted separately to determine their percentage in relation to the total mass of the whole seed.

**Figure 1.**
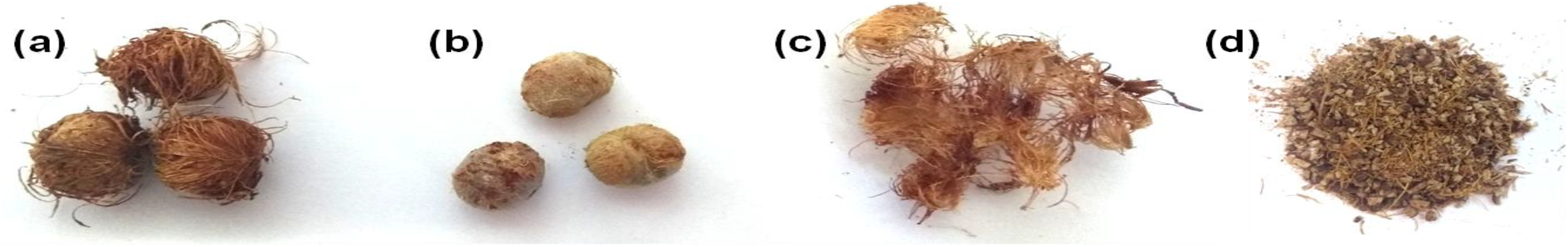
Açaí seed samples: (a) whole seeds; (b) core stone after removing the external fibers; (c) fiber layer; (d) milled whole seeds.

#### 2.2.1 Determination of extractives, carbohydrates, and acid insoluble solids

*In natura* milled açaí seeds underwent an extraction process (Sluiter et al., 2008) with some modifications. Approximately 2 g of the biomass were weighted into cellulose thimbles and extracted with water, which was followed by a 95% ethanol extraction; each extraction step was performed for at least 12 h. The procedure was carried out using six Soxhlet apparatus in parallel. By the end of the extraction, three of the thimbles were put in a 105 ºC drying oven overnight to calculate the extractives by weight difference, while the other three were put in a 40 ºC drying oven to be used in the following chemical characterization step. Then, 0.3 g of the dried, extractive-free *in natura* and acid-treated açaí seed samples were submitted to an acid hydrolysis process in two steps in triplicate (Sluiter et al., 2012). In the first step, the samples were mixed with 3 mL of a 72% sulfuric acid solution in round-bottom hydrolysis tubes and put in a 30 °C water bath for 1 h under constant stirring. In the second step, 84 mL of deionized water were added to the tubes, which were autoclaved for 1 h at 121 °C. After this, the solutions were vacuum filtered through dried, preweighted Gooch crucibles. The acidic liquors were neutralized with CaCO_3_ and went through HPLC and HPAEC-PAD analysis, which is described below, for carbohydrate quantification. The crucibles containing the remaining solids were dried overnight in an oven at 105 ºC, and the dry weight was recorded for acid insoluble solids (AIS) quantification using the difference in weight. The insoluble ash content was also measured using the difference in weight after the same crucibles were put overnight in a furnace at 575 ºC.

#### 2.2.2. X-ray diffraction analysis

X-ray diffraction (XRD) analyses of the milled açaí seed samples were performed using Bruker's D8 Advance equipment, Cu Kα radiation ("lambda" = 1.5418 angstroms), 40 kVA voltage and 40 mA current, a scanning angle in the range of 2 ≤ 2θ ≤ 60 degrees, and acquisition time of 0.6 s per step.

### 2.3. Diluted-acid hydrolysis of açaí seeds for mannose release

Four sulfuric acid concentrations were evaluated for the diluted-acid hydrolysis step, corresponding to 1.5%, 3.0%, 3.5%, and 4.5% (% w/w). Each of these solutions was evaluated at a 30-and a 60-min residence time at 121 °C. Each condition was performed in at least four replicates in round-bottom hydrolysis tubes containing 4 g (dry weight) of the milled açaí seeds and 16 mL of the corresponding diluted-acid solution, resulting in a solid:liquid ratio of 1:4. The tubes were put in an autoclave for 30-or 60-min at 121 ºC and then cooled in an ice bath. After this, 64 mL of water were added to the tubes, which were agitated for homogenization, and samples of the liquid streams were withdrawn, being then filtrated, neutralized, and prepared for HPLC and HPAEC-PAD analysis, to determine the sugar and degradation products, as described below.

The solid contents of two of the four tubes were filtrated in preweighted fiber glass filters and put in an oven at 105 ºC overnight to calculate the amount of mass transferred to the acidic liquid phase. The solid contents of the other two tubes were filtered and stored in the refrigerator until further use either for characterization of the chemical composition or for enzymatic hydrolysis assays. Prior to the characterization assays, the samples were dried at 40 °C until reaching less than 10% moisture and then used for the determination of AIS, ash, and carbohydrates, as previously described.

The combined severity factor was calculated for each diluted-acid hydrolysis condition, which was evaluated based on the severity factor R_0_, which accounts for the effect of the temperature, residence time, and pH of the hydrolysates after the reaction, through the expression Log R_0_-pH, where Log R_0_ is given by equation [1](Ferreira-Leitão et al., 2010):

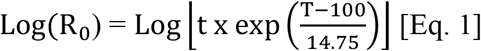

where *t* is the reaction time of the pretreatment in minutes, and *T* is the reaction temperature in °C.

### 2.4. Enzyme activity measurements and enzymatic hydrolysis assays

The endomannanase activity of mannanase BGM “Amano” 10 was determined using a 0.5% locust bean gum (Sigma-Aldrich) solution as the substrate. The enzyme solution was diluted in a 50 mM sodium citrate buffer (pH 4.8), and an aliquot of 0.25 mL was mixed with 0.25 mL of the substrate solution and incubated in a water bath for 30 min at 50 ºC. Then, 0.5 mL of 3,5-dinitrosalicylic acid (DNS), prepared according to Teixeira et al. (Teixeira et al., 2012), was added to each tube after 30 min of incubation to stop the reaction, and the tubes were put in a boiling water bath for 5 min. The absorbance of the colored solutions was measured via spectrophotometer (ThermoScientific Evolution 201) at a wavelength of 540 nm to quantify the reducing sugars. One unit of endomannanase was defined as the amount of enzyme required to release 1 µmol of reducing sugars equivalent to mannose in 1 min at 50 °C.

β-mannosidase activity was determined by adding 100 μL of 10 mM of 4-nitrophenyl β-*D*-mannopyranoside to 200 μL of a 0.5 M sodium citrate buffer (pH 4.8), 600 μL of distilled water, and 100 μL of an appropriately diluted enzyme sample. The assay was incubated at 50 °C for 10 min, and the reaction was stopped with the addition of 500 μL of 1.0 M sodium carbonate. The liberation of *p*-nitrophenol was monitored via spectrophotometer at a wavelength of 405 nm. One unit of β-mannanase was defined as the amount of enzyme that released 1 μmol of *p*-nitrophenol for 1 min at 50 °C.

The enzymatic hydrolysis assays were performed in 50-mL flasks, with a total assay mass of 20 g containing 2–20% (w/w) of biomass (native seeds and seed samples after the acid hydrolysis) based on its dry weight, the enzyme solution in a 0.05 M sodium citrate buffer (pH 4.8), and 0.02% sodium azide. The amount of enzyme added was such that the mannanase activity load was 400 UI per gram of biomass, which was established in the preliminary assays. The flasks were incubated in a shaker at 50 °C and 200 rpm. Aliquots were withdrawn at 0, 2, 6, 24, 48, and 72 h and analyzed by HPLC for sugar quantification. Mannose yields were calculated according to equation [2].

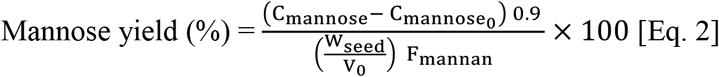

where C_mannose_ is the mannose concentration in the hydrolysates (g/L); C_mannose_0__ is the initial mannose concentration in the hydrolysis assay; W_seed_ is the total weight of the seed in the hydrolysis assay (g); V_0_ is the initial volume of the liquid (L); F_mannan_ is the initial mass fraction of mannan in samples.

### 2.5. Chromatographic conditions

Sugars and acetic acid were quantified by HPLC using an Ultimate 3000 system (Thermo Scientific, USA) equipped with a refractive index detector RI-101 (SHODEX, Japan). For sugar quantification, an Aminex HPX-87P (300 x 7.8 mm, Bio-Rad) column was used, with a Carbo-P precolumn (Bio-Rad, USA) and an inline deashing system (Bio-Rad). The mobile phase used ultrapure water at a flow rate of 0.6 mL/min with an oven temperature of 80 °C and detector temperature of 60 °C. The sugar composition of açaí seed samples and of acidic and enzymatic hydrolysates were also cross-checked by monosaccharides and disaccharides identification and quantification using a Thermo Scientific Dionex ICS-5000 system (Canada) using high-performance anion exchange chromatography with pulse amperometric detection (HPAEC-PAD). The guard cartridge and analytical column used were the CarboPac PA1 (Thermo Scientific, 4 mm x 50 mm and 10 µm particle sizes) and CarboPac PA1 (Thermo Scientific, 4 mm x 250 mm and 10 µm particle sizes). The column temperature was 15 °C, and the mobile phase was composed of phase A (type 1 reagent-grade deionized water) and phase B (300 mM NaOH solution). The gradient programs used for the separation were as follows: 0.0–32.0 min, 0% B; 32.0–32.1 min, 0–85% B; 32.1–42.0 min, 85% B; 42.0–42.1 min, 85–0% B; and 42.1–52.0 min, 0% B. The flow rate was 1.25 mL/min, and the injection volume was 5 µL. The system was also equipped with a postcolumn addition of 450 mM NaOH solution with a flow rate of 0.8 mL/min.

The acetic acid was quantified using an Aminex HPX-87H (300 x 7.8 mm, Bio-Rad) column with a Carbo-H precolumn (Bio-Rad, USA) and an inline deashing system (Bio-Rad). The mobile phase was 5 mM H_2_SO_4_ at a flow rate of 0.6 mL/min with oven and detector temperatures of 30 °C and 45 °C, respectively. Furfural, hydroxymethylfurfural, and phenolic compounds (gallic acid, hydroxybenzoic acid, vanillin, ferulic acid, and cinnamic acid) were quantified with the diode array detector DAD-3000 (Thermo Scientific, USA). The column was a LiChroCART RP-18e (4.6 × 250 mm, Merck, Germany) equipped with the precolumn LiChroCART RP-18e (4.0 × 4.0 mm, Merck, Germany). The mobile phase was composed of phase A (type 1 reagent-grade deionized water) and phase B (methanol) at a flow rate of 0.4 mL/min, with oven temperatures of 30 °C and detector wavelengths of 280 and 320 nm. The gradient programs used for separation were as follows: 0.0–3.1 min, 15% B; 3.1–8.1 min, 65% B; 8.1–8.2 min, 95% B; 8.1–9.9 min, 95% B; 9.9–14.0 min, 0% B; 14.0–19.0 min, 15% B. Concentrations were quantified by external calibration.

## 3. Results and Discussion

### 3.1. Açaí seed chemical characterization

First, 35 samples of dried mature açaí seed samples were weighted to determine the proportion of the mass of the external fiber layer to the whole seed samples (Figure 1). By botanical definition, the external fiber layer is not considered part of the seed; however, we denominated the seed as the residue generated after the depulping and sieving of açaí berries (fibers + seed) because—for the sake of brevity—it is improbable that any large-scale commercial use of this residue will separate those fractions.

The average weight of the whole seeds was 0.78 g ± 0.12, ranging from 0.56 g to 1.06 g, and the mass percentage of fiber in relation to the whole seed was equivalent to 5.97% ± 1.45. These data are in close agreement with a previous study that reported that the whole seeds average weight was 0.72 ± 0.04 g and that the fibrous layer corresponded to 6.50% of the whole seed weight (Pessoa et al., 2010).

The literature data regarding the açaí seed composition so far is conflicting. Therefore, to better evaluate the seed’s uses, a confirmation of its chemical composition is crucial to design the most suitable processing methods for sugar recovery. In the current study, the characterization of two distinct seed samples lots was performed, as well as an analysis of different seed fractions. Table 1 presents the composition of the milled samples of whole seed from the two lots and of the core stones and fiber layer of one lot. The composition of the whole seed showed that the material is mostly composed of carbohydrates, with mannan—a polymer of mannose— being its main component and corresponding to 47.09% and 52.46% of its total dry weight for lot 1 and lot 2, respectively. Smaller amounts of other structural sugars were also identified, such as glucose, xylose, galactose, and arabinose. Rambo et al. (2015) reported a similar composition for açaí seeds, corresponding to 53.6% mannose, 8.66% glucose, 3.18% xylose, 1.43% galactose, 0.69% arabinose, and 0.17% rhamnose. The seed’s lipid content was not analyzed in the current study; however, it has been reported that *Euterpe oleracea* seeds contain only 0.33% total fat (Wycoff et al., 2015).

**Table 1.** Chemical composition of the whole açaí seeds from two different lots and of the core stone and fiber layer of one lot, here expressed as a percentage of dry matter.

| Component | Dry mass (%) |  |  |  |
| --- | --- | --- | --- | --- |
|  | whole seed<br>(lot 1) | Core stone<br>(lot 1) | Fiber layer<br>(lot 1) | whole seed<sup>a</sup><br>(lot 2) |
| Anhydromannose | 47.09 ± 1.42 | 47.19 ± 2.58 | nd<sup>c</sup> | 52.46 ± 1.51 |
| Anhydroglucose | 6.09 ± 0.67 | 4.61 ± 0.48 | 21.88 ± 0.46 | 8.40 ± 0.52 |
| Anhydroxylose | 1.83 ± 0.33 | 1.13 ± 0.16 | 15.12 ± 0.39 | 2.05 ± 0.22 |
| Anhydrogalactose | 1.79 ± 0.21 | 2.61 ± 0.12 | 0.90 ± 0.04 | 1.51 ± 0.27 |
| Anhydroarabinose | 0.40 ± 0.02 | 0.85 ± 0.03 | 0.82 ± 0.03 | 0.63 ± 0.03 |
| AIS<sup>b</sup> | 18.34 ± 0.64 | 18.36 ± 0.61 | 31.80 ± 0.36 | 19.54 ± 1.56 |
| Extractives | 15.45 ± 0.95 | 16.72 ± 2.43 | 12.89 ± 1.88 | 9.89 ± 2.09 |
| Ash | 0.61 ± 0.09 | 0.41 ± 0.03 | 2.12 ± 0.06 | 0.44 ± 0.02 |

<sup>a</sup> The characterization of the core stones and fibers are shown only for lot 1 because lot 2 was received milled from the producer. <sup>b</sup> AIS: acid insoluble solids account for the organic matter that was insoluble after acid hydrolysis condition and is calculated by not counting the amount of acid insoluble ash. <sup>c</sup> nd: not detected.

The composition of the core stones (Fig. 1b) showed a high similarity with the whole seed, as expected, considering that this fraction corresponds to almost 94% of the whole seed’s mass. The fibers, however, presented a distinct sugar profile with no detectable mannose content and higher contents of glucan, xylan, and AIS when compared with the core stone. AIS can be presumably counted as lignin; however, because açaí seeds are quite different from typical lignocellulosic biomass, further analyses of this AIS are required to confirm if all of its content corresponds to the lignin. The high percentage of extractives in açaí seeds is in accordance with its reported polyphenolic polymeric procyanidins content (Melo et al., 2016). Nevertheless, it is possible that not all the content of the polymeric procyanidins is accounted for in the extractives because hydrogen bonds can be formed between the hydroxyl groups of polyphenols and oxygen of the glycosidic linkages of polysaccharides, making procyanidins imprisoned in the cell wall of carbohydrates and not extractable using organic solvents (Jakobek, 2015).

The high mannan content confirmed in the current study contradicts most studies reporting on açaí seeds’ composition, which have stated cellulose as the main polysaccharide in the seed (Altman, 1956; Oliveira et al., 2015, 2013; Rodríguez-Zúñiga et al., 2008; Wycoff et al., 2015). Although Rambo et al. (2015) quantified the carbohydrate content of açaí seeds and reported mannan as the main polysaccharide, no further discussion was made about this finding in that study. The fact that many studies reported high cellulose content in the seed could be related to the use of indirect methods to determine the material’s composition, which measure the total fiber content instead of specific sugar quantification using chromatographic methods. For example, Altman (1956) employed a method developed by Waksman and Stevens (1930) which is based on the principle that in general, hemicellulose is hydrolyzed upon treatment with diluted acid, while cellulose and lignin are resistant. However, because the mannan from the açaí seeds is—much like cellulose—highly recalcitrant toward diluted-acid hydrolysis, mannan has been incorrectly quantified as cellulose. Wycoff et al. (2015) have analyzed açaí seed samples by nuclear magnetic resonance and observed peaks related to glycosidic bonds, inferring that these were related to cellulose and hemicellulose and citing previous studies that indicate cellulose as the seed’s main polysaccharide.

Cellulose has an unusual crystallinity among biopolymers, and XRD is the most commonly used technique to obtain data for cellulose crystallinity. Therefore, XRD analyses were performed to verify if the açaí seed samples had the typical diffraction profile of materials containing high amounts of cellulose. Figure 2 presents the XRD profiles of the milled samples of the whole açaí seeds (stone plus fibers), the açaí seed stone, and the fibers.

**Figure 2.**
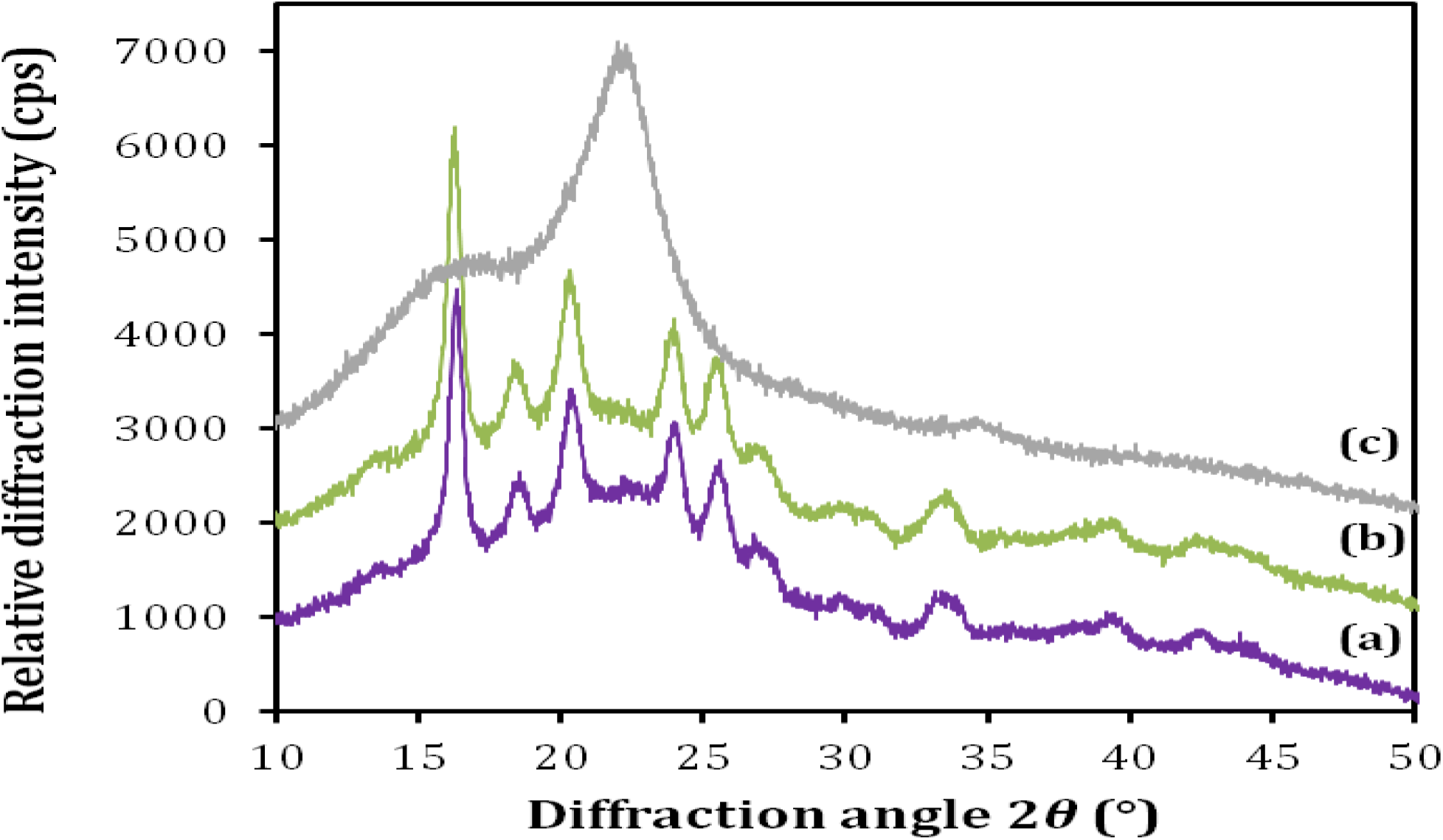
XRD profiles of the milled samples of (a) whole açaí seed; (b) açaí seed core stone, and (c) açaí seed fibers.

The açaí seed fiber presented a typical cellulose I diffraction pattern with two peaks at 2*θ* equal to 16.0 and 22.0, which correspond to the cellulose crystal planes 110 and 200, respectively (French and Santiago Cintrón, 2013). The fibers’ XRD profile was very similar to the ones found for other agricultural residues, such as sugarcane bagasse and wheat straw, which are biomasses that contain ~40% cellulose (da Silva et al., 2010). These data are in accordance with the macroscopic aspect of the fibrous material, as well as with its high glucan content. However, the samples of milled whole seeds or açaí stones did not show a diffraction peak corresponding to the cellulose crystalline plane 200. A comparison of the diffraction patterns, which were completely different in the 15º–25º region, which is characteristic of cellulose crystals, also corroborates the sugar composition data for each fraction analyzed, confirming that açaí seeds do not contain cellulose as their main polysaccharide.

This high mannose content in açaí seeds is supported by the fact that the secondary walls of the endosperm cells in the seeds of many species contain very little cellulose (Bento et al., 2013); these consist of noncellulosic cell wall storage polysaccharides that are digested during germination, being usually mannans, galactomannans, glucomannans, xyloglucans, and galactans (Aspinall, 1959; Moreira and Filho, 2008; Scheller and Ulvskov, 2010). A number of botanical studies have identified mannan as a customary reserve polysaccharide in seeds from the Arecaceae (palm trees) family (Ishrud et al., 2001; Mazzottini-dos-Santos et al., 2017; Neto et al., 2010). However, these studies have focused on the aspects of plant physiology and seed germination and have not quantitatively evaluated the seed’s polysaccharide chemical composition. Even though mannans can be common cell wall storage polysaccharides, it is interesting that the açaí seed’s mannan content of about 50% of its dry weight is quite high. Most vegetal biomasses, such as grasses and hardwoods, have low amounts of mannose (<2%), while softwood can reach up to 15% (Hu et al., 2016). Palm kernel cakes, copra meal, and spent coffee grounds are agro-industrial residues with the highest reported mannose content, reaching 30-35%, 28-32%, and 14-19% of their dry weight, respectively (Cerveró et al., 2010; Fan et al., 2014; Khuwijitjaru et al., 2012; Kusakabe et al., 1987; Nguyen et al., 2019; Pedras et al., 2019).

The reported chemical structure and composition of other mannose-rich seeds or residues indicate that the most common polysaccharides in these materials are β-1,4-mannan and/or galactomannan, which has a (1→4)-β-D-mannopyranosyl backbone and can alternatively be substituted by α-D-galactopyranosyl residues at position *O*-6 (Bento et al., 2013; Moreira and Filho, 2008). For example, the endosperm of many seeds from small leguminous trees, as well as the locust bean and guar gums, have galactomannan as the main polysaccharide, with a galactose:mannose molar ratio of 1:2 to 1:4 (Moreira and Filho, 2008). In contrast, the mature palm kernel, coconut copra meal, ivory nuts, and green coffee beans have mannans composed of linear chains of 1,4-linked β-D-mannopyranosyl residues that contain less than 5% galactose and small amounts of other polysaccharides (Aspinall, 1959). Therefore, considering the monosaccharide’s profile obtained from the acid hydrolysis of açaí seeds (Table 1) and the data from the literature for other seeds, it is hypothesized that a linear β-1,4-mannan is the main polysaccharide of the mature açaí seed. The very low content of galactose in mature açaí seeds renders the presence of galactomannan unlikely because this polysaccharide is reported to have a galactose:mannose molar ratio of 1:2. However, further studies to elucidate the carbohydrate structure will be necessary.

### 3.2. Effect of acid hydrolysis for mannose release from açaí seeds

Moderate diluted-acid hydrolysis was evaluated as a possible strategy to release mannose from açaí seeds through the breakdown of the mannan, making the monomeric sugars readily available in the acid’s liquid phase. Acid hydrolysis was evaluated at a fixed temperature of 121 °C and by varying the H_2_SO_4_ concentration and residence time. Table 2 presents the severity factor for each condition evaluated, the percentage of insoluble solids recovered, and its chemical composition and the sugar composition of the hydrolysates obtained.

**Table 2.** Characterization of the recovered insoluble solids and sugar composition of the hydrolysates from the acid hydrolysis of açaí seeds at different H_2_SO_4_ concentrations and residence times.

| % H<sub>2</sub>SO<sub>4</sub><br>(m/m) | Time<br>(min) | % RIS<sup>a</sup> | R<sub>0</sub><sup>b</sup> | Content in recovered solid (%) |  |  |  | Sugar concentration in the hydrolysate (g/L) |  |  |  |  |
| --- | --- | --- | --- | --- | --- | --- | --- | --- | --- | --- | --- | --- |
|  |  |  |  | Mannan | Glucan | Xylan | AIS<sup>c</sup> | Man | Gal | Xyl | Ara | Glu |
| Un<sup>d</sup> | - | 100 | - | 52.46 ± 1.51 | 8.40 ± 0.52 | 2.05 ± 0.22 | 19.54 ± 1.56 | - | - | - | - | - |
| 1.5% | 30 | 81.1 ± 0.1 | 0.99 | 60.39 ± 1.67 | 10.01 ± 0.10 | 2.46 ± 0.20 | 23.72 ± 0.94 | 13.54 ± 2.10 | 2.12 ± 0.22 | 1.03 ± 0.16 | 1.58 ± 0.03 | 0.51 ± 0.01 |
| 3.0% | 30 | 73.5 ± 0.4 | 1.19 | 57.32 ± 0.72 | 8.99 ± 1.21 | 1.38 ± 0.19 | 22.39 ± 0.88 | 23.28 ± 1.65 | 2.71 ± 0.12 | 1.88 ± 0.07 | 1.38 ± 0.02 | 0.67 ± 0.03 |
| 3.5% | 30 | 71.6 ± 0.1 | 1.32 | 55.80 ± 1.32 | 9.72 ± 0.15 | 0.83 ± 0.09 | 27.84 ± 1.38 | 27.65 ± 0.37 | 3.01 ± 0.08 | 2.21 ± 0.01 | 1.41 ± 0.06 | 0.78 ± 0.08 |
| 4.5% | 30 | 71.6 ± 0.7 | 1.29 | 53.36 ± 1.12 | 10.13 ± 0.21 | 1.05 ± 0.04 | 29.22 ± 1.54 | 29.93 ± 1.83 | 2.98 ± 0.18 | 2.48 ± 0.33 | 1.31 ± 0.08 | 0.84 ± 0.02 |
| 1.5% | 60 | 72.8 ± 0.5 | 1.30 | 58.99 ± 0.99 | 9.66 ± 0.05 | 1.49 ± 0.10 | 27.28 ± 1.43 | 27.12 ± 0.13 | 3.14 ± 0.03 | 2.57 ± 0.13 | 1.68 ± 0.02 | 0.87 ± 0.01 |
| 3.0% | 60 | 70.2 ± 0.2 | 1.44 | 54.10 ± 2.20 | 10.28 ± 0.76 | 0.98 ± 0.10 | 27.07 ± 0.57 | 41.76 ± 1.09 | 3.55 ± 0.05 | 3.45 ± 0.24 | 1.50 ± 0.03 | 1.07 ± 0.06 |
| 3.5% | 60 | 70.4 ± 0.6 | 1.64 | 51.34 ± 3.10 | 10.32 ± 0.56 | 0.81 ± 0.14 | 25.05 ± 2.28 | 46.22 ± 0.44 | 3.58 ± 0.06 | 3.26 ± 0.21 | 1.48 ± 0.03 | 1.14 ± 0.11 |
| 4.5% | 60 | 62.3 ± 0.9 | 1.70 | 55.25 ± 1.18 | 10.42 ± 0.40 | 0.40 ± 0.07 | 30.86 ± 3.09 | 49.54 ± 0.20 | 3.38 ± 0.02 | 3.10 ± 0.34 | 1.39 ± 0.02 | 1.34 ± 0.12 |

<sup>a</sup>RIS: recovered insoluble solids; <sup>b</sup>R<sub>0</sub>: combined severity factor; <sup>c</sup>AIS: acid insoluble solids; <sup>d</sup>Un: untreated açai seed from lot 2.

From Table 2, there is a correlation between the severity of the acid hydrolysis, the percentage of insoluble solids recovered, and the concentration of mannose released, indicating that to a certain extent, more biomass components are transferred into the liquid phase when the severity is higher, which was expected. The lowest severity condition (R_0_ 0.99) resulted in 81.1% recovered solids, while for the most severe condition, this value decreased to 62.3%. The duration of the treatment, from 30–60 min, had an important impact on the release of sugars. Mannose concentration increased according to the increase in acid concentration from 1.5% to 4.5% H_2_SO_4_, ranging from 13.54 g/L to 29.93 g/L (9.4% to 20.7% yield) for hydrolysis carried out for 30 min and from 27.12 g/L to 49.54 g/L (18.8% to 34.4% yield) when treatments were carried out for 60 min. The same pattern could be observed for glucose. Similarly, the diluted-acid hydrolysis of nondilapidated spent coffee grounds—a residue rich in mannan—resulted in 85–70% solids recovery after hydrolysis, with acid concentrations ranging from 1–5% v/v and residence times from 30–60 min at 95 °C (Juarez et al., 2018).

Even though mannose and glucose concentrations increased with a higher severity, the xylose, arabinose and galactose concentrations in the hydrolysates decreased during the most severe conditions, indicating the partial degradation of these sugars in the liquid fraction. This is in accordance with studies that reported a lower activation energy for the hydrolysis of xylan (101 kJ/mol) than for mannan (113 kJ/mol) and cellobiose (110 kJ/mol) hydrolysis (Canettieri et al., 2007; Mosier et al., 2002; Nattorp et al., 1999). It is well known that the combination of high temperature, acidic pH, and prolonged reaction time may contribute to the formation of undesired compounds derived from sugar dehydration, such as furfural and hydroxymethylfurfural, as well as the degradation of phenolic structures (Ferreira-Leitão et al., 2010; Oral et al., 2014). Therefore, to evaluate acid hydrolysis conditions, one should take into account both the sugar release yield and formation of degradation products. Moreover, it is also important to calculate the total mannose recovery because higher severity treatments may result in a higher release of mannose, but also in a partial degradation of this sugar, causing an overall loss of the desired product. In the present study, low amounts of hydroxymethyl furfural were quantified in the hydrolysates and were equivalent to 56 mg/L, which was detected only in the most severe condition (4.5% H_2_SO_4_, 60 min). Very low concentrations of acetic acid were quantified in the hydrolysates from all conditions, which were in the range of 60–210 mg/L. Other compounds, such as furfural, vanillin, gallic, ferulic, cinnamic, and hydrobenzoic acids were monitored but were either under the limit of quantification or not detected. Figure 3 shows the mannose recovery balance in both the solid and liquid fractions obtained after the acid hydrolysis of seeds for each condition. Biomass characterization protocols are multistep and very laborious procedures, but overall, over 95% of the original mannose content could be detected either in the hydrolysate or preserved in the solid fraction, which is in good agreement with the absence/low concentration of the degradation products detected (Figure 3). The high mannose recovery indicates that at the temperature evaluated, the acid concentration and reaction duration were at an acceptable range for mannose stability.

**Figure 3.**
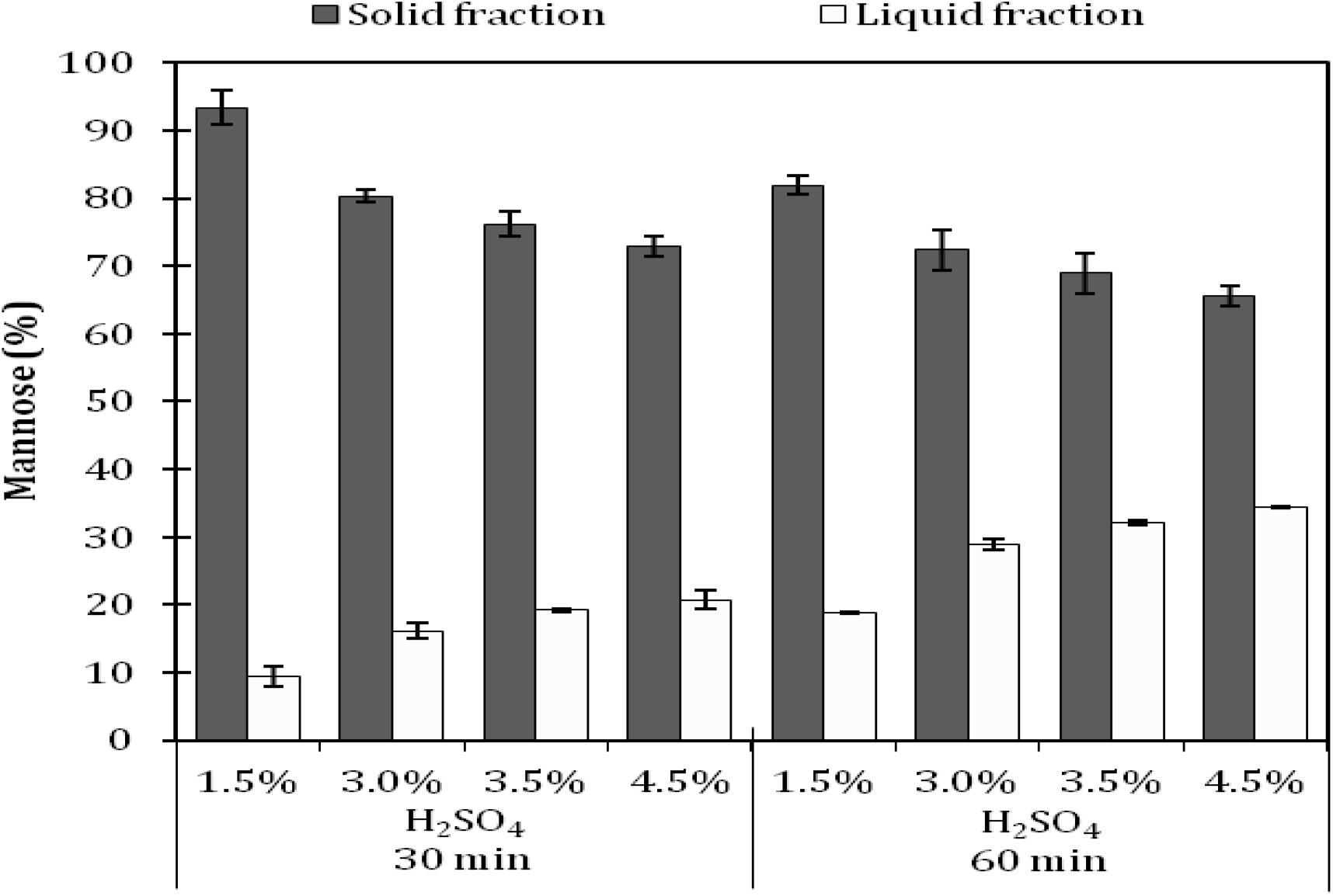
Percentage of mannose recovery from the milled açaí seeds after acid hydrolysis with diluted H_2_SO_4_ (1.5%–4.5%) for 30 and 60 min of retention time at 121 °C. White bars: Percentage of mannose recovered in the liquid fraction after acid hydrolysis. Dark gray bars: mannose content retained in the solid fraction after acid hydrolysis.

Optimized diluted-acid hydrolysis of xylans at mild temperatures can reach yields of 80–95%, while cellulose is much more resistant to diluted acid attacks at temperatures under 200 °C (Wyman et al., 2004). Açaí seed mannan’s susceptibility to diluted-acid hydrolysis seems to lie in between because 34.3% of the mannan could be converted into mannose during the most severe condition evaluated. Reducing sugars when in solution exist as a mix of cyclic structures, which are more resistant to degradation, and as an open chain, which is the more reactive acyclic carbonyl form (Angyal, 1984). In polysaccharides, the sugar at the chain end or ramified end will exist in ring and open chain structures, being more susceptible to degradation (Nattorp et al., 1999). Additionally, it has been shown that at room temperature, glucose and mannose present a lower carbonyl percentage than galactose, xylose, and arabinose (Hayward and J. Angyal, 1977). These data correlate to the low concentration of the sugar degradation products found in the hydrolysates and to the fact that water-insoluble and linear mannans, which are likely cellulose, can form crystalline structures (Chanzy et al., 1987, 1984) that are recalcitrant and resistant to diluted sulfuric acid attack; however, other amorphous hemicelluloses, such as galactoarabinoxylans, are known to be highly susceptible to diluted-acid hydrolysis (Wyman et al., 2004).

The current knowledge of the chemical hydrolysis of mannan is limited because there are only a few reports of mannose production from vegetal biomass through acid hydrolysis. One study (Fan et al., 2014) evaluated the diluted sulfuric acid hydrolysis under microwave irradiation of deproteinated palm kernel cake (PKC) containing 55.71% mannan; the results of that study showed a mannose yield of 92% at the optimized condition of 148 °C, 0.75 N H_2_SO_4_ (equivalent to 3.5% w/w), 10.5 min, and a solid:liquid (S:L) ratio of 1:50. Besides the use of microwave irradiation assistance, higher temperatures and the possible differences between deproteinated PKC and açaí seed recalcitrance to acid attack, the higher mannose yields achieved for PKC acid hydrolysis may have been favored by the low substrate concentration evaluated (S:L ratio of 1:50). In the present study, a S:L of 1:4 was used because a low substrate concentration leads to extremely diluted hydrolysates that are not feasible for industrial applications.

The mild conditions for the acid hydrolysis of mannan that were evaluated were not sufficient to efficiently break down the recalcitrance of this polysaccharide because 65–93% of the original mannose content remained in the solids recovered from the seed’s diluted-acid hydrolysis (Figure 3). However, increasing the process’ severity could result in the high formation of degradation products along with the generation of very toxic and corrosive streams, although there are different approaches for the removal of degradation products from the hydrolysate (Mussatto and Roberto, 2004). Nevertheless, in this study it was decided to avoid their formation during hydrolysis. Therefore, a sequential process of enzymatic hydrolysis with the mannanases of the recovered solids was evaluated to attempt to further release the mannose.

### 3.3. Enzymatic hydrolysis for mannose release from recovered solids from acid hydrolysis

After a preliminary screening of several commercial and lab-made enzyme preparations, the enzyme mannanase BGM “Amano” 10 (Amano Enzyme Inc., Japan) was selected as the most efficient for the hydrolysis of açaí seed’s mannan. The enzyme preparation, which is commercially available as a powder, had activities of β-mannanase and β-mannosidase of 26,750 IU/g and 15.05 IU/g, respectively. Figure 4 shows the mannose yields obtained with the enzymatic hydrolysis of the acid-treated seed samples compared with the native milled seeds.

**Figure 4.**
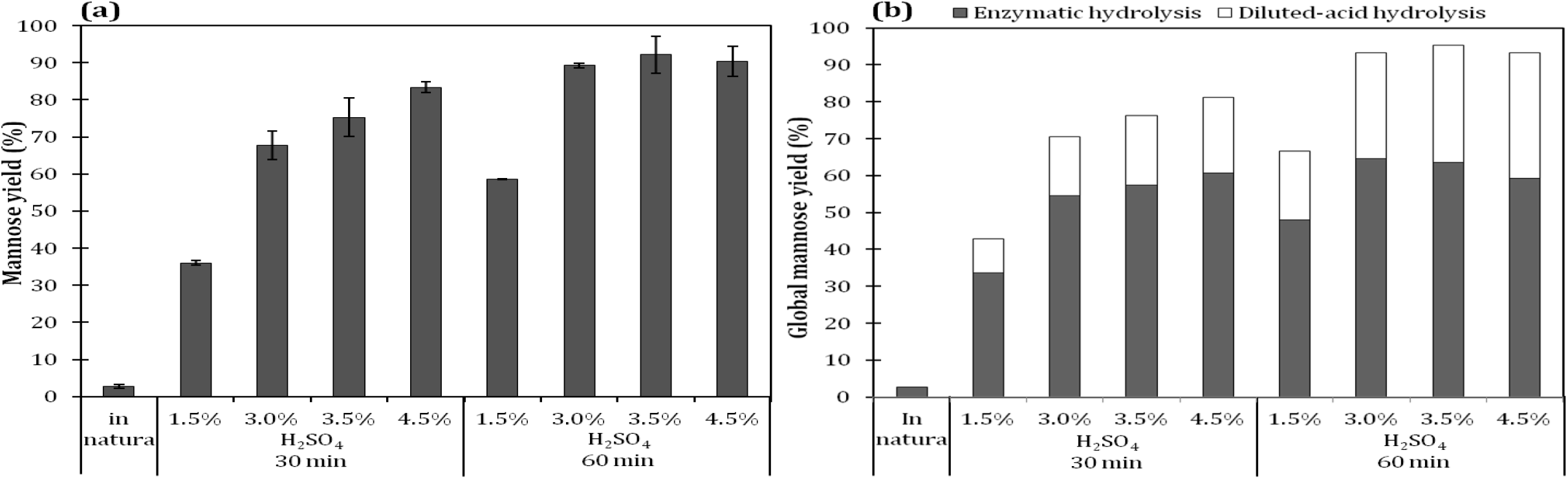
(a) Mannose yield obtained after 72 h of enzymatic hydrolysis of *in natura* and previously acid-hydrolyzed açaí seed milled samples; (b) Global mannose yield calculated in relation to the initial mannose content in the native seed. The assays were conducted with 2% solids and 400 UI of mannanase BGM “Amano” 10 per gram of sample.

The native açaí seed sample was highly recalcitrant to the enzymatic attack, resulting in a less than 3% mannose yield. Nonetheless, after the material had been partially digested by sulfuric acid, it became much more susceptible to the attack of the mannanases, resulting in a 90% mannose yield for samples that were treated for 60 min with 3%, 3.5%, and 4.5% sulfuric acid. Consequently, the recovery of mannose could be substantially increased through a sequential process of diluted-acid hydrolysis followed by enzymatic hydrolysis, potentially reaching over 93% global mannose recovery for the most favorable conditions when using both the acid and enzymatic hydrolysis steps (Figure 4b).

Data regarding the enzymatic hydrolysis of mannan into mannose is scarce in the literature because there are not many abundant agroindustrial residues rich in this polysaccharide. So far, there have been no studies in the literature exploring mannose production from açaí seeds, but there are some reports with other residues, such as PKC, copra meal, and spent coffee grounds. A study that evaluated the enzymatic hydrolysis of PKC, which contains 35.2% of mannan, reported that this residue was readily hydrolyzed into mannose with a mixture of two enzymes (Mannaway and Gammanase) with no previous treatment, resulting in an 87% conversion rate of mannan into mannose after 96 h (Cerveró et al., 2010). Most of mannan in PKC consists of a (1→4)-linked mannan with a low degree of substitution with galactose (Düsterhöft et al., 1991), which is the same structure that we hypothesized for the mannan in açaí seeds. However, in the present study, native açaí seeds were poorly hydrolyzed by mannanases, reaching only 3% conversion of mannan to mannose, suggesting that these residues have distinctive mannan structures and/or the mannan is less accessible in açaí seeds because of interactions with other structural and nonstructural components.

The set of experiments presented in Figure 4 were performed with a 2% açaí seed content (w/w), which led to high yield but also to hydrolysates containing low concentrations of mannose of about 11 g/L of at the best conditions. To have an effective industrial process, it is of the utmost importance to work on concentrated media to reduce the capital cost of equipment and the use of water. Therefore, the effect of solids loading on the enzymatic hydrolysis was evaluated in a range of 2–20% (Figure 5). Samples treated with 3% acid for 60 min were selected for the assays because the seeds treated with 3%, 3.5%, and 4.5% of sulfuric acid were equally susceptible to mannanase attack (Figure 4), and this condition has a lower impact on the use of H_2_SO_4_, formation of acidic effluents, and degradation products.

**Figure 5.**
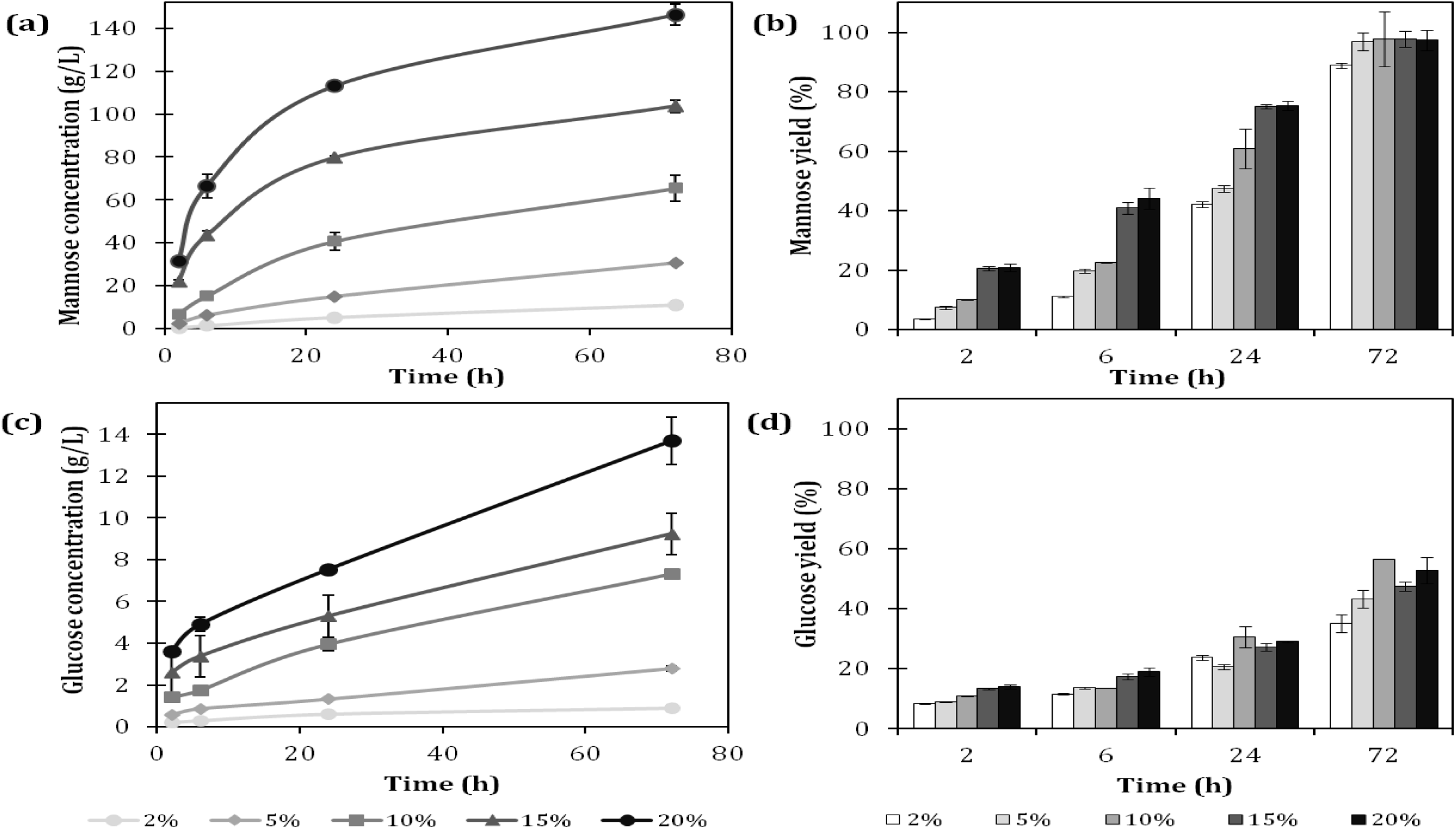
Enzymatic hydrolysis profile at the different solid contents of acid-hydrolyzed açaí seeds. (a) Mannose concentration; (b) mannose yield; (c) glucose concentration; (d) glucose yield. The assays were conducted with 400 UI of mannanase BGM “Amano” 10 per gram of sample. The samples were priory treated with 3% H_2_SO_4_ for 60 min at 121 °C.

The enzymatic hydrolysis in the assays containing 2%, 5%, 10%, 15%, and 20% of acid-treated açaí seeds resulted in mannose concentrations, respectively, of 9.8, 30.6, 65.4, 103.7, and 146.3 g/L after 72 h. Regardless of the solid content evaluated (from 5–20%), the conversion of mannan content into mannose reached over 95% (Figure 5b). The high yields achieved indicate that the endomannanase and β-mannosidase balance in the commercial enzymatic preparation used is adequate for the complete hydrolysis of açaí seed’s mannan. To the best of our knowledge, the mannose concentration reached in the assays with 20% solids is by far the highest reported in the literature for the enzymatic hydrolysis of an agricultural residue.

Only mannose and glucose were detected in the hydrolysates by HPLC analysis. At 72 h, the glucose concentrations reached 0.9, 2.8, 7.3, 9.2, and 13.7 g/L for the assays containing 2%, 5%, 10%, 15%, and 20% of acid-hydrolyzed açaí seed, respectively. It is very interesting to note that roughly, a glucose:mannose ratio of 1:10 could be observed (See Suplementary Material). Considering that the enzyme used is a mannanase with nearly no cellulase activity, we hypothesize that the glucose released during enzymatic hydrolysis is derived from the mannan structure. The glucose:mannose ratio of 1:10 derived from mannan hydrolysis is in agreement with the definition of a “true” mannan, which relates to polysaccharides with more than 85– 95% mannose content and a high degree of uniformity in the structure (Aspinall, 1959; Stephen, 1983).

The mannose and glucose concentrations obtained at 72 h of hydrolysis showed a linear relationship to the initial solid content, indicating that no significative inhibition effect took place during mannan hydrolysis. These data differ from what is typically reported for the enzymatic hydrolysis of cellulosic substrates because it has been shown that by increasing the substrate concentration, the corresponding yield decreases (Kristensen et al., 2009). Although the 72 h mannose yields reached a plateau for all the solids content evaluated, the mannan conversion rate was faster in hydrolysis assays with higher solid contents, which also has an opposite effect to what is observed in the hydrolysis of fibrous cellulose-rich materials. This fact reinforces the observation that açaí seed mannan hydrolysis occurred in a pattern that differs greatly from the enzymatic hydrolysis of lignocellulosic materials by cellulases, which are affected by the “solids effect” including substrate effects, product inhibition, water content constraints, enzyme adsorption characteristics, and others (Modenbach and Nokes, 2013).

A similar evaluation was performed for the enzymatic hydrolysis of PKC using different substrate concentrations from 5–20% (w/v), reaching, at optimized conditions, a mannose concentration of 67.5 g/L. In agreement with our observation, Shukor et al. (2016), reported a direct increase in the production of simple sugars with an increase of the PKC content, which indicated that no substrate inhibition effect was taking place. However, the authors did not present a hydrolysis profile over time or the PKC characterized, which restricts other comparisons with the present study.

A recent study has shown that 15 g of mannose could be obtained for every 100 g of spent coffee ground (SCG) after the removal of the non-saccharide content after delignification and defatting of SCG (Nguyen et al., 2019). At the present study, 57.5 g of mannose could be obtained for every 100 g of *in natura* açaí seed, with a total mannose recovery of 98.6%. Figure 6 presents the mass balance of the overall process for the mannose release from açaí seeds considering the sequential process of diluted acid hydrolysis and an enzymatic hydrolysis step with 20% solids. The results presented in the current study demonstrate that mannan from açaí seeds could be a low-cost source to produce mannose in high yields and concentrations. The development of this field could fill the present market demand for the cost-effective production of mannose and its derivatives (Hu et al., 2016), which is hindered by the scarce sources of mannose and development of appropriate methods.

**Figure 6.**
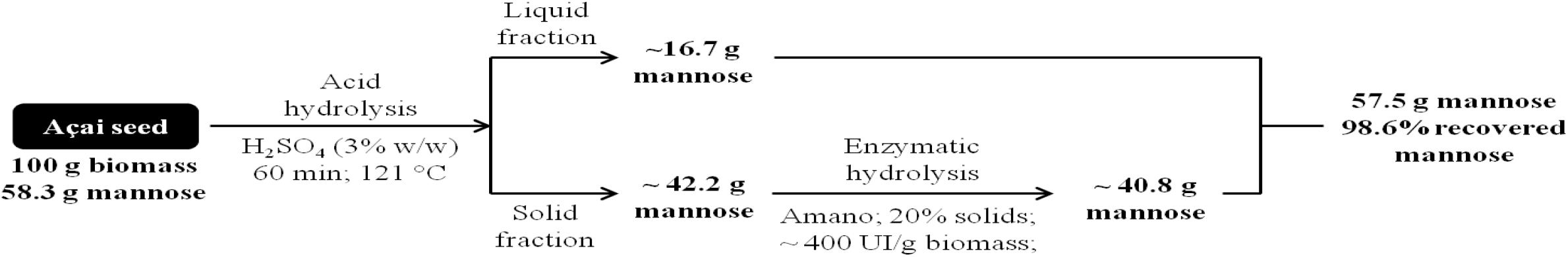
Mass balance of mannose production from milled açaí seeds treated with 3% H_2_SO_4_ for 60 min at 121 °C followed by enzymatic hydrolysis with BGM Amano 10 and 20% solid content.

## 4. Conclusions

In this pioneer study, a sequential process of diluted-acid and enzymatic hydrolysis of açaí seeds was developed to convert its high mannan content into mannose. Mannanases-catalyzed hydrolysis of acid-treated seeds resulted in 146 g/L of mannose and a 96.8% yield. To the best of our knowledge, this is by far the highest concentration of mannose reported for the enzymatic hydrolysis of an agricultural residue, which could open new perspectives for mannose use as a platform molecule. Finally, giving a proper destination to açaí seeds could add value to the whole açaí productive chain while promoting development in the Amazon region.

## Appendix. Supplementary data

E-supplementary data of this work can be found in online version of the paper.

## Conflicts of interest

All authors are listed as inventors of a patent application (BR 10 2018 067282 7) that includes the process described in this paper.

## Acknowledgment

This work was supported by the Serrapilheira Institute (grant number Serra-1708-15009) and the Coordination for the Improvement of Higher Education Personnel (CAPES – grant AUXPE 0415/2016). We want to thank M.Sc Leandro Barbosa and the Bioethanol Laboratory at the Federal University of Rio de Janeiro for their assistance with the HPAEC-PAD analysis and the Laboratory of Combinatorial Catalysis at the National Institute of Technology for the XRD measurements. Açaí Amazonas Ltd. and Amano Enzyme Inc. are thanked for providing açaí seeds and enzyme samples, respectively. M.Sc. Ronaldo Rodrigues is acknowledged for his assistance with the graphical abstract. Ingrid Santos Miguez and João Silva were granted scholarships, respectively, from Fundação Carlos Chagas Filho de Amparo à Pesquisa do Estado do Rio de Janeiro (FAPERJ) and the National Council for Scientific and Technological Development (PIBITI/CNPq).

## References

Altman, R.F.A., 1956. Estudo químico de plantas amazônicas. Boletim Técnico do Instituto Agronômico do Norte 31, 109–111.

Angyal, S.J., 1984. The composition of reducing sugars in solution. Adv. Carbohydr. Chem. Biochem. 42, 15–68. https://doi.org/10.1016/S0065-2318(08)60122-5

Ariandi, Yopi, Meryandini, A., 2015. Enzymatic hydrolysis of copra meal by mannanase from Streptomyces sp. BF3.1 for the production of mannooligosaccharides. J. Biosci. 22, 79–86. https://doi.org/10.4308/hjb.22.2.79

Aspinall, G.O., 1959. Structural chemistry of the hemicelluloses. Adv. Carbohydr. Chem. 14, 429–468. https://doi.org/10.1016/S0096-5332(08)60228-3

Bento, J.F., Mazzaro, I., Silva, L.M.A., Moreira, R.A., Ferreira, M.L.C., Reicher, F., Petkowicz, C.L.O., 2013. Diverse patterns of cell wall mannan/galactomannan occurrence in seeds of the Leguminosae. Carbohydr. Polym. 92, 192–199. https://doi.org/10.1016/j.carbpol.2012.08.113

Canettieri, E.V., Rocha, G.J.M., de Carvalho Jr, J.A., de Almeida e Silva, J.B., 2007. Optimization of acid hydrolysis from the hemicellulosic fraction of Eucalyptus grandis residue using response surface methodology. Bioresour. Technol. 98, 422–428. https://doi.org/10.1016/j.biortech.2005.12.012

Cerveró, J.M., Skovgaard, P.A., Felby, C., Sørensen, H.R., Jørgensen, H., 2010. Enzymatic hydrolysis and fermentation of palm kernel press cake for production of bioethanol. Enzyme Microb. Technol. 46, 177–184. https://doi.org/10.1016/j.enzmictec.2009.10.012

Chanzy, H., Perez, S., Miller, D.P., Paradossi, G., Winter, W.T., 1987. An electron diffraction study of the mannan I crystal and molecular structure. Macromolecules 20, 2407–2413. https://doi.org/10.1021/ma00176a014

Chanzy, H.D., Grosrenaud, A., Vuong, R., Mackie, W., 1984. The crystalline polymorphism of mannan in plant cell walls and after recrystallisation. Planta 161, 320–329. https://doi.org/10.1007/BF00398722

da Silva, A.S., Inoue, H., Endo, T., Yano, S., Bon, E.P.S., 2010. Milling pretreatment of sugarcane bagasse and straw for enzymatic hydrolysis and ethanol fermentation. Bioresour. Technol. 101. https://doi.org/10.1016/j.biortech.2010.05.008

Düsterhöft, E., Voragen, A.G.J., Engels, F.M., 1991. Non‐starch polysaccharides from sunflower (Helianthus annuus) meal and palm kernel (Elaeis guineenis) meal— preparation of cell wall material and extraction of polysaccharide fractions. J. Sci. Food Agric. 55, 411–422. https://doi.org/10.1002/jsfa.2740550309

Fan, S.P., Jiang, L.Q., Chia, C.H., Fang, Z., Zakaria, S., Chee, K.L., 2014. High yield production of sugars from deproteinated palm kernel cake under microwave irradiation via dilute sulfuric acid hydrolysis. Bioresour. Technol. 153, 69–78. https://doi.org/10.1016/j.biortech.2013.11.055

Ferreira-Leitão, V., Perrone, C.C., Rodrigues, J., Franke, A.P.M., Macrelli, S., Zacchi, G., 2010. An approach to the utilisation of CO_2_ as impregnating agent in steam pretreatment of sugar cane bagasse and leaves for ethanol production. Biotechnol. Biofuels 3, 1–8. https://doi.org/10.1186/1754-6834-3-7

Fioravanti, C., 2013. Açaí: Do pé para o lanche. Pesqui. FAPESP 64–68. URL http://revistapesquisa.fapesp.br/2013/01/11/acai-do-pe-para-o-lanche/

French, A.D., Cintrón, M.S., 2013. Cellulose polymorphy, crystallite size, and the segal crystallinity index. Cellulose 20, 583–588. https://doi.org/10.1007/s10570-012-9833-y

Hayward, L.D.J., Angyal, S., 1977. A symmetry rule for the circular dichroism of reducing sugars, and the proportion of carbonyl forms in aqueous solutions thereof. Carbohydr. Res. 53, 13–20. https://doi.org/10.1016/S0008-6215(00)85450–6

Hu, X., Shi, Y., Zhang, P., Miao, M., Zhang, T., Jiang, B., 2016. d-Mannose: Properties, production, and applications: An overview. Compr. Rev. Food Sci. Food Saf. 15, 773–785. https://doi.org/10.1111/1541-4337.12211

Ishrud, O., Zahid, M., Zhou, H., Pan, Y., 2001. A water-soluble galactomannan from the seeds of Phoenix dactylifera L. Carbohydr. Res. 335, 297–301. https://doi.org/10.1016/S0008-6215(01)00245-2

Jakobek, L., 2015. Interactions of polyphenols with carbohydrates, lipids and proteins. Food Chem. 175, 556–567. https://doi.org/10.1016/j.foodchem.2014.12.013

Jiang, M., Li, H., Shi, J., Xu, Z., 2018. Depolymerized konjac glucomannan: preparation and application in health care. J. Zhejiang Univ. B 19, 505–514. https://doi.org/10.1631/jzus.B1700310

Juarez, G.F.Y., Pabiloña, K.B.C., Manlangit, K.B.L., Go, A.W., 2018. Direct dilute acid hydrolysis of spent coffee grounds: A new approach in sugar and lipid recovery. Waste and Biomass Valorization 9, 235–246. https://doi.org/10.1007/s12649-016-9813-9

Khuwijitjaru, P., Watsanit, K., Adachi, S., 2012. Carbohydrate content and composition of product from subcritical water treatment of coconut meal. J. Ind. Eng. Chem. 18, 225–229. https://doi.org/10.1016/j.jiec.2011.11.010

Kristensen, J.B., Felby, C., Jørgensen, H., 2009. Yield-determining factors in high-solids enzymatic hydrolysis of lignocellulose. Biotechnol. Biofuels 2, 1–10. https://doi.org/10.1186/1754-6834-2-11

Kusakabe, I., Zama, M., Park, G.G., Tubaki, K., Murakami, K., 1987. Preparation of β-1, 4-Mannobiose from white copra meal by a mannanase from penicillium purpurogenum. Agric. Biol. Chem. 51, 2825–2826. https://doi.org/10.1080/00021369.1987.10868480

Lim, W.S., Lee, J.W., 2013. Influence of pretreatment condition on the fermentable sugar production and enzymatic hydrolysis of dilute acid-pretreated mixed softwood. Bioresour. Technol. 140, 306–311. https://doi.org/10.1016/j.biortech.2013.04.103

Makkee, M., Kieboom, A.P.G., Bekkum, H. van, 1985. Production methods of D-mannitol. Starch/Stärke 37, 136–141.

Mazzottini-dos-Santos, H.C., Ribeiro, L.M., Oliveira, D.M.T., 2017. Roles of the haustorium and endosperm during the development of seedlings of Acrocomia aculeata (Arecaceae): dynamics of reserve mobilization and accumulation. Protoplasma 254, 1563–1578. https://doi.org/10.1007/s00709-016-1048-x

Melo, P.S., Arrivetti, L.O.R., de Alencar, S.M., Skibsted, L.H., 2016. Antioxidative and prooxidative effects in food lipids and synergism with α-tocopherol of açaí seed extracts and grape rachis extracts. Food Chem. 213, 440–449. https://doi.org/10.1016/j.foodchem.2016.06.101

Mishra, D.K., Hwang, J.S., 2013. Selective hydrogenation of d-mannose to d-mannitol using NiO-modified TiO_2_ (NiO-TiO2) supported ruthenium catalyst. Appl. Catal. A Gen. 453, 13–19. https://doi.org/10.1016/j.apcata.2012.11.042

Modenbach, A.A., Nokes, S.E., 2013. Enzymatic hydrolysis of biomass at high-solids loadings - A review. Biomass and Bioenergy 56, 526–544. https://doi.org/10.1016/j.biombioe.2013.05.031 Review

Moreira, L.R.S., Filho, E.X.F., 2008. An overview of mannan structure and mannan-degrading enzyme systems. Appl. Microbiol. Biotechnol. 79, 165–178. https://doi.org/10.1007/s00253-008-1423-4

Mosier, N.S., Ladisch, C.M., Ladisch, M.R., 2002. Characterization of acid catalytic domains for cellulose hydrolysis and glucose degradation. Biotechnol. Bioeng. 79, 610–618. https://doi.org/10.1002/bit.10316

Mussatto, S.I., Roberto, I.C., 2004. Alternatives for detoxification of diluted-acid lignocellulosic hydrolyzates for use in fermentative processes: A review. Bioresour. Technol. 93, 1–10. https://doi.org/10.1016/j.biortech.2003.10.005

Nattorp, A., Graf, M., Spühler, C., Renken, A., 1999. Model for random hydrolysis and end degradation of linear polysaccharides: Application to the thermal treatment of mannan in solution. Ind. Eng. Chem. Res. 38, 2919–2926. https://doi.org/10.1021/ie990034j

Neto, M.A.M., Lobato, A.K.D.S., Alves, J.D., Goulart, P.D.F.P., Laughinghouse IV, H.D., 2010. Seed and seedling anatomy in Euterpe oleracea Mart. during the germination process. J. Food, Agric. Environ. 8, 1147–1152.

Nguyen, Q.A., Cho, E.J., Lee, D.S., Bae, H.J., 2019. Development of an advanced integrative process to create valuable biosugars including manno-oligosaccharides and mannose from spent coffee grounds. Bioresour. Technol. 272, 209–216. https://doi.org/10.1016/j.biortech.2018.10.018

Nogueira, A.K.M., de Santana, A.C., Garcia, W.S., 2013. A dinâmica do mercado de açaí fruto no Estado do Pará: De 1994 a 2009. Rev. Ceres 60, 324–331. https://doi.org/10.1590/S0034-737X2013000300004

Oliveira, J.A.R., Komesu, A., Maciel Filho, R., 2013. Hydrothermal pretreatment for enhancing enzymatic hydrolysis of seeds of açaí (Euterpe oleracea) and sugar recover. Chem. Eng. Trans. 64, 2656–2663. https://doi.org/10.3969/j.issn.0438-1157.2013.07.047

Oliveira, J.A.R., Martins, L.H.S., Komesu, A., Maciel Filho, R., 2015. Evaluation of alkaline delignification (NaOH) of açaí seeds (eutherpe oleracea) treated with H_2_SO_4_ dilute and effect on enzymatic hydrolysis. Chem. Eng. Trans. 43, 499–504. https://doi.org/10.3303/CET1543084

Oral, R.A., Mortas, M., Dogan, M., Sarioglu, K., Yazici, F., 2014. New approaches to determination of HMF. Food Chem. 143, 367–370. https://doi.org/10.1016/j.foodchem.2013.07.135

Otieno, D.O., Ahring, B.K., 2012. The potential for oligosaccharide production from the hemicellulose fraction of biomasses through pretreatment processes: Xylooligosaccharides (XOS), arabinooligosaccharides (AOS), and mannooligosaccharides (MOS). Carbohydr. Res. 360, 84–92. https://doi.org/10.1016/j.carres.2012.07.017

Pedras, B.M., Nascimento, M., Sá-Nogueira, I., Simões, P., Paiva, A., Barreiros, S., 2019. Semi-continuous extraction/hydrolysis of spent coffee grounds with subcritical water. J. Ind. Eng. Chem. 8–11. https://doi.org/10.1016/j.jiec.2019.01.001

Pessoa, J.D.C., Arduin, M., Martins, M.A., de Carvalho, J.E.U., 2010. Characterization of Açaí (E. oleracea) fruits and its processing residues. Brazilian Arch. Biol. Technol. 53, 1451–1460. https://doi.org/10.1590/S1516-89132010000600022

Pompeu, D.R., Silva, E.M., Rogez, H., 2009. Optimisation of the solvent extraction of phenolic antioxidants from fruits of Euterpe oleracea using response surface methodology. Bioresour. Technol. 100, 6076–6082. https://doi.org/10.1016/j.biortech.2009.03.083

Rambo, M.K.D., Schmidt, F.L., Ferreira, M.M.C., 2015. Analysis of the lignocellulosic components of biomass residues for biorefinery opportunities. Talanta 144, 696–703. https://doi.org/10.1016/j.talanta.2015.06.045

Rodríguez-Zúñiga, U.F., Lemo, V., Farinas, C.S., Neto, V.B., Couri, S., 2008. Evaluation of agroindustrial residues as substrates for cellulolytic enzymes production under solid state fermentation, in: 7th Brazilian MRS Meeting.

Rungrassamee, W., Kingcha, Y., Srimarut, Y., Maibunkaew, S., Karoonuthaisiri, N., Visessanguan, W., 2014. Mannooligosaccharides from copra meal improves survival of the Pacific white shrimp (Litopenaeus vannamei) after exposure to Vibrio harveyi. Aquaculture 434, 403–410. https://doi.org/10.1016/j.aquaculture.2014.08.032

Scheller, H.V., Ulvskov, P., 2010. Hemicelluloses. Annu. Rev. Plant Biol. 61, 263–289. https://doi.org/10.1146/annurev-arplant-042809-112315

Shukor, H., Abdeshahian, P., Al-Shorgani, N.K.N., Hamid, A.A., Rahman, N.A., Kalil, M.S., 2016. Enhanced mannan-derived fermentable sugars of palm kernel cake by mannanase-catalyzed hydrolysis for production of biobutanol. Bioresour. Technol. 218, 257–264. https://doi.org/10.1016/j.biortech.2016.06.084

Sluiter, A., Hames, B., Ruiz, R.O., Scarlata, C., Sluiter, J., Templeton, D., Crocker, D., 2012. Determination of structural carbohydrates and lignin in biomass, National Renewable Energy Laboratory Analytical Procedure.

Sluiter, A., Ruiz, R., Scarlata, C., Sluiter, J., Templeton, D., 2008. Determination of extractives in biomass, National Renewable Energy Laboratory Analytical Procedure.

Srivastava, P.K., Kapoor, M., 2017. Production, properties, and applications of endo-β-mannanases. Biotechnol. Adv. 35, 1–19. https://doi.org/10.1016/j.biotechadv.2016.11.001

Stephen, A.M., 1983. Other Plant Polysaccharides, in: The Polysaccharides. Elsevier, pp. 97–193. https://doi.org/10.1016/B978-0-12-065602-8.50008-X

Teixeira, R.S.S., Da Silva, A.S.A., Ferreira-Leitão, V.S., Da Silva Bon, E.P., 2012. Amino acids interference on the quantification of reducing sugars by the 3,5-dinitrosalicylic acid assay mislead carbohydrase activity measurements. Carbohydr. Res. 363, 33–37. https://doi.org/10.1016/j.carres.2012.09.024

van Zyl, W.H., Rose, S.H., Trollope, K., Görgens, J.F., 2010. Fungal β-mannanases: Mannan hydrolysis, heterologous production and biotechnological applications. Process Biochem. 45, 1203–1213. https://doi.org/10.1016/j.procbio.2010.05.011

Waksman, A., Stevens, S.R., 1930. A system of proximate chemical analysis of plant materials. Ind. Eng. Chem.

Wycoff, W., Luo, R., Schauss, A.G., Neal-Kababick, J., Sabaa-Srur, A.U.O., Maia, J.G.S., Tran, K., Richards, K.M., Smith, R.E., 2015. Chemical and nutritional analysis of seeds from purple and white açaí (Euterpe oleracea Mart.). J. Food Compos. Anal. 41, 181–187. https://doi.org/10.1016/j.jfca.2015.01.021

Wyman, C.E., Decker, S.R., Himmel, M.E., Brady, J.W., Skopec, C.E., Viikari, L., 2004. Hydrolysis of cellulose and hemicellulose, in: Dumitriu, S. (Ed.), Polysaccharides: Structural diversity and functional versatility. Marcel Dekker, Inc., New York, pp. 995–1033.

Yamaguchi, K.K.D.L., Pereira, L.F.R., Lamarão, C.V., Lima, E.S., da Veiga-Junior, V.F., 2015. Amazon acai: Chemistry and biological activities: A review. Food Chem. 179, 137–151. https://doi.org/10.1016/j.foodchem.2015.01.055

