## Supplemental Table 1 and Figure 1 for "High concentration and yield production of mannose from açaí (*Euterpe oleracea*) seeds via diluted-acid and mannanase-catalyzed hydrolysis"

### Supplementary Material

**Supplementary Table 1.** Sugar concentration (g/L) released during the enzymatic hydrolysis with different solids loading, and the respective mannose/glucose ratio.

| Solids loading (%) | Sugar concentration (g/L) | | Man/Glu Ratio |
|---|---|---|---|
| | Mannose | Glucose | |
| 20 | 146.3 | 13.7 | 10.7 |
| 15 | 103.7 | 9.2 | 11.3 |
| 10 | 65.4 | 7.3 | 9.0 |
| 5 | 30.6 | 2.8 | 10.9 |
| 2 | 10.9 | 0.9 | 12.1 |

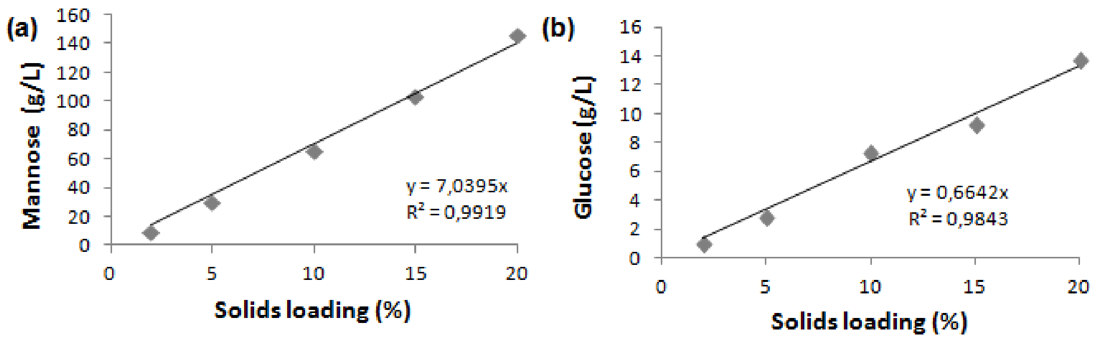


**Supplementary Figure 1.** (a) Mannose and (b) glucose concentration (g/L) released during the enzymatic hydrolysis versus solids loading in the assay.
